# FORGE-KI: A Modular Framework for Endogenous Knock-In Engineering Across HDR and PITCh/MMEJ Repair Pathways

**DOI:** 10.64898/2026.06.15.732404

**Authors:** Dylan Conklin, Ji-Ann Lee, Michael Palazzolo, Steven M. Dubinett, Jay M. Lee

## Abstract

Targeted knock-in technologies have enabled precise insertion of reporters, affinity tags, degrons, and other functional payloads into endogenous genomic loci. Over the past decade, a diverse collection of genome engineering strategies has emerged, including approaches based on homology-directed repair (HDR), microhomology-mediated end joining (MMEJ), homology-mediated end joining (HMEJ), and related methodologies. While these advances have greatly expanded the capabilities of endogenous genome engineering, they have also increased the complexity of donor design, assembly, and validation.

Here, we describe FORGE-KI (Functional Oncology Research Genetic Engineering – Knock in), a pathway-matched design workflow for endogenous knock-in engineering that aligns the assembly strategy with the underlying repair mechanism. For large-cargo insertions, we use a modular five-component framework that separates gene-specific targeting arms from reusable functional modules, allowing rapid assembly of HDR donor constructs targeting AHR, IRF1, and FOSL1 from a shared reagent collection. For MMEJ/PITCh applications, where short targeting elements permit rapid fabrication, we developed a streamlined one-step pipeline in which the entire donor and selection payload is synthesized as a single continuous fragment for direct cloning, compressing the design-to-reagent cycle time. This MMEJ workflow is paired with a dual-promoter nuclease vector (pForge-KI-MMEJ-Cas9-DualGuide) that drives the PITCh-release and locus-specific guides from distinct promoters, a design intended to reduce the repeated-promoter instability associated with some dual-guide vectors.

We also established a standardized workflow for donor assembly, generation of knock-in cell populations, molecular validation, and selectable-cassette removal, and we demonstrate it by generating a functional, selection-marker-free, cytokine-inducible IRF1 HDR reporter line and an inducible IRF1 PITCh/MMEJ reporter pool with confirmed junction enrichment. In parallel, we developed forgeKI, an R package that automates C-terminal reporter knock-in design across both HDR and PITCh/MMEJ repair pathways, including guide selection, target-biology validation, targeting-arm design, domestication, donor-assembly planning, and generation of synthesis-ready constructs.

Together, the reagents and software provide a practical system for endogenous knock-in engineering that supports multiple payloads, selection strategies, and repair pathways within a shared donor organization. Rather than replacing existing knock-in technologies, this framework provides a modular foundation for incorporating, extending, and automating the published knock-in methods.

## Introduction

The development of programmable nucleases has transformed the ability to engineer endogenous genomic loci^1^. Over the past decade, a diverse collection of knock-in strategies has emerged, including approaches based on homology-directed repair (HDR), microhomology-mediated end joining (MMEJ), homology-mediated end joining (HMEJ)^2^, HITI^3^, GeneWeld^4^, and related methodologies. Collectively, these technologies have demonstrated that precise insertion of reporters, affinity tags, degrons, selectable markers, and other functional payloads can be achieved across a wide range of biological systems.

As the field has matured, the challenge has increasingly shifted from demonstrating targeted integration to managing the growing complexity of donor design. Modern knock-in experiments often require selection among multiple repair strategies, reporter formats, selection schemes, and validation approaches^5^. The optimal donor architecture depends strongly on the biological question being addressed. For example, HiBiT fusions provide highly sensitive protein detection^6^, fluorescent reporters enable flow-cytometric and imaging-based analyses, and HaloTag- or degron-based payloads support chemical biology and targeted protein degradation applications^7^. Each configuration offers distinct advantages, but also introduces additional design and cloning requirements.

Similar considerations apply to selection strategies. Many successful knock-in approaches rely on reporters or resistance genes expressed directly from the endogenous locus. While this design is elegant and effective for highly expressed genes, low endogenous expression can limit the utility of reporter- or resistance-based selection. To address this challenge, numerous groups have adopted removable selection cassettes driven by independent promoters. These cassettes facilitate enrichment of correctly engineered cells while allowing removal of the selection module following establishment of the knock-in allele^8^. In addition, cell lines frequently contain pre-existing fluorescent proteins and antibiotic-resistance markers from prior engineering efforts, creating a practical need for alternative reporter and selection configurations.

Importantly, the objective of the present work was not to identify a single optimal knock-in architecture. Different experimental goals frequently require different reporter payloads, selection strategies, and donor configurations. We therefore sought to develop a framework that could accommodate multiple best-practice design solutions rather than prescribe a single implementation, supporting variation and extensibility while preserving a common organizational structure for donor construction^9,10^.

Donor backbone design presents additional considerations. Random donor integration remains an important practical concern, motivating incorporation of counterselection strategies that improve recovery of correctly modified cells^11,12^. At the same time, the growing number of available payloads, selectable markers, repair pathways, and target loci creates an increasingly large design space, so the effort required to design, assemble, and validate knock-in reagents often scales more rapidly than the underlying genome engineering technologies themselves.

We therefore sought to develop an operational framework that aligns the reagent-generation strategy with the constraints of the chosen repair pathway. For HDR, where kilobase-scale targeting arms make single-fragment synthesis of the complete donor impractical, we describe a modular framework that separates gene-specific elements from reusable functional components across a standardized five-component architecture. For MMEJ/PITCh pathways, the minute size of the required microhomology sequences allows the entire donor cassette to be synthesized as a single continuous fragment, enabling a streamlined one-step direct-cloning pipeline. To complement this split workflow, we introduce a dual-promoter nuclease vector (pForge-KI-MMEJ-Cas9-DualGuide). Together, this hybrid approach preserves structural flexibility, reduces the repeated-promoter instability associated with some dual-guide designs, and matches each genome engineering strategy to its most efficient assembly pathway.

## Results

### Design of a modular framework for endogenous knock-in engineering

A wide range of genome engineering strategies support targeted gene insertion, including HDR, MMEJ, HMEJ, and related approaches. Although these methods differ in donor architecture and repair mechanism, many share common functional requirements, including reporter payloads, selectable markers, and vector backbones. We reasoned that separating gene-specific elements from reusable functional components could simplify donor construction while maintaining compatibility with multiple repair pathways.

To achieve this, we developed a modular knock-in architecture consisting of five functional components: a 5′ targeting arm, a fusion module, a selectable cassette, a 3′ targeting arm, and an acceptor backbone (Figure 1). Gene-specific information is isolated within the 5′ and 3′ targeting arms, while reporter payloads, selectable cassettes, and backbone elements are maintained as reusable modules. This organization enables donor constructs to be assembled from a small set of standardized parts while preserving flexibility in target locus, reporter configuration, and repair strategy.

**Figure 1.**
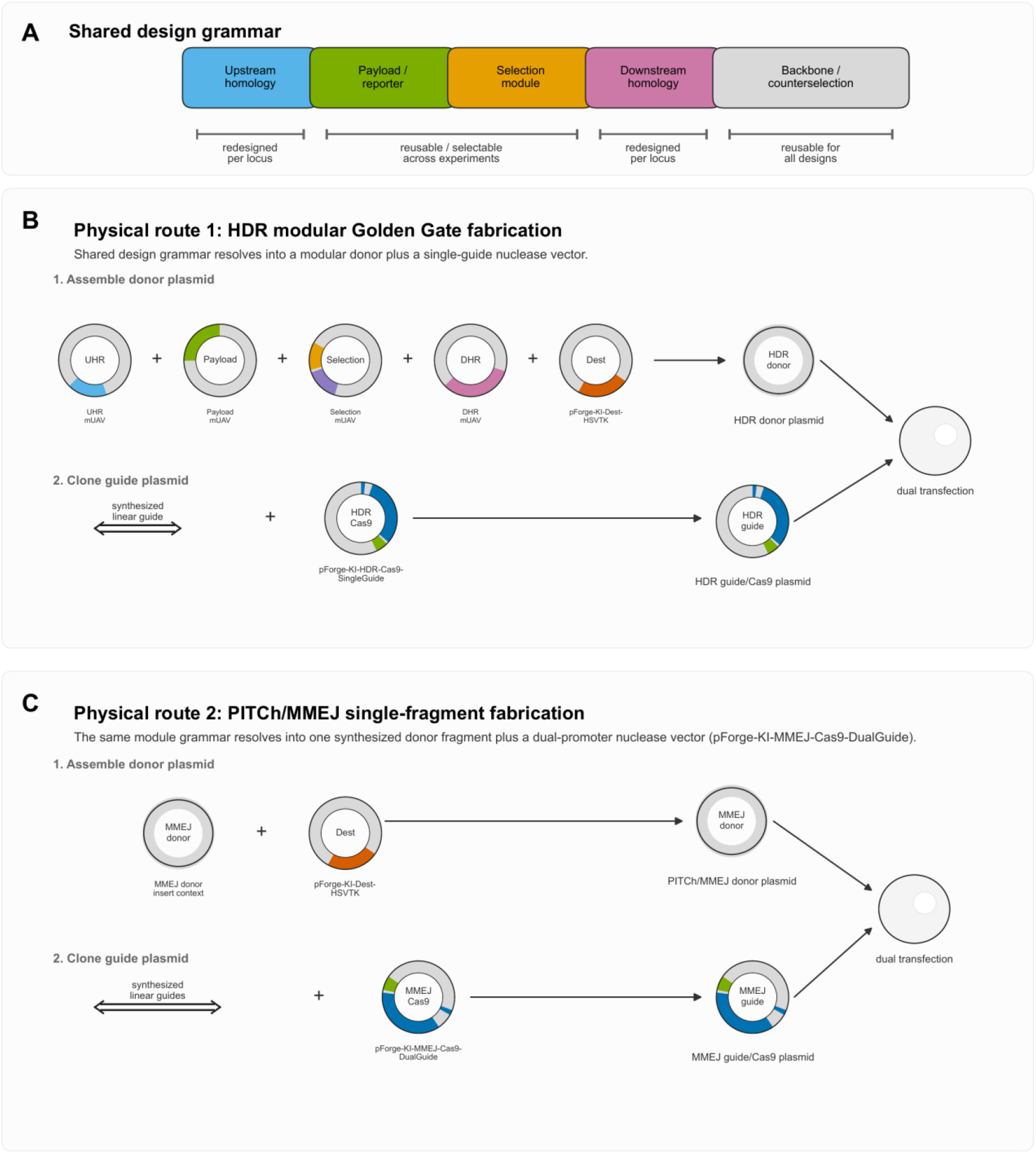
Modular architecture and dual-pathway assembly for endogenous knock-in engineering. (A) Shared five-module donor organization: locus-specific 5′ and 3′ homology arms (redesigned per locus) flank reusable payload/reporter, selection, and backbone/counterselection modules. (B) HDR route: the donor plasmid is assembled from five modules by one-pot BsaI Golden Gate into the pForge-KI-Dest-HSVTK acceptor, the locus-specific guide is cloned into a single-guide Cas9 vector, and the two plasmids are co-transfected. (C) PITCh/MMEJ route: the entire donor is synthesized as a single fragment and cloned in one step into a counterselection-compatible KanR/ori backbone (PGK–HSV-TK outside the integrating cassette; yielding p1538), while the locus and PITCh donor-release guides are supplied by the dual-guide nuclease vector (pForge-KI-MMEJ-Cas9-DualGuide), and the two plasmids are co-transfected.

We matched the cloning methodology to the structural requirements of each repair mechanism. While the functional payloads remain conceptually consistent, their physical assembly pathways diverge based on targeting-arm size. For HDR applications, the kilobase-scale homology arms necessitate a multi-part modular assembly using the mUAV entry system to build the donor step by step into the pForge-KI-Dest-HSVTK backbone. For PITCh/MMEJ applications, the small size of the microhomology sequences enables the entire donor cassette to be synthesized as a single continuous fragment and cloned in a single step into a counterselection-compatible KanR/ori backbone carrying a PGK–HSV-TK marker outside the integrating cassette^13^. This hybrid approach provides separate fabrication routes for both short- and long-homology strategies^14^.

### A modular reagent collection for endogenous knock-in engineering

To support rapid construction of endogenous knock-in donors, we established a modular reagent collection consisting of reusable fusion modules, selectable cassettes, and a common acceptor backbone (Figure 2). A single acceptor backbone (pForge-KI-Dest-HSVTK) was developed containing an HSV-TK counterselection cassette. Reusable fusion modules included HiBiT, GFP11^15^, ddDegron^16^, dTAG^17^ (FKBP12^F36V), Halo-HiBiT^7^, P2A-EGFP, and HiBiT-P2A-EGFP. Three selectable-cassette modules were generated (mRFP1-Hygro, mRFP1-Puro, and BFP-Puro). These modules were maintained independently and combined with gene-specific homology regions through Golden Gate assembly.

**Figure 2.**
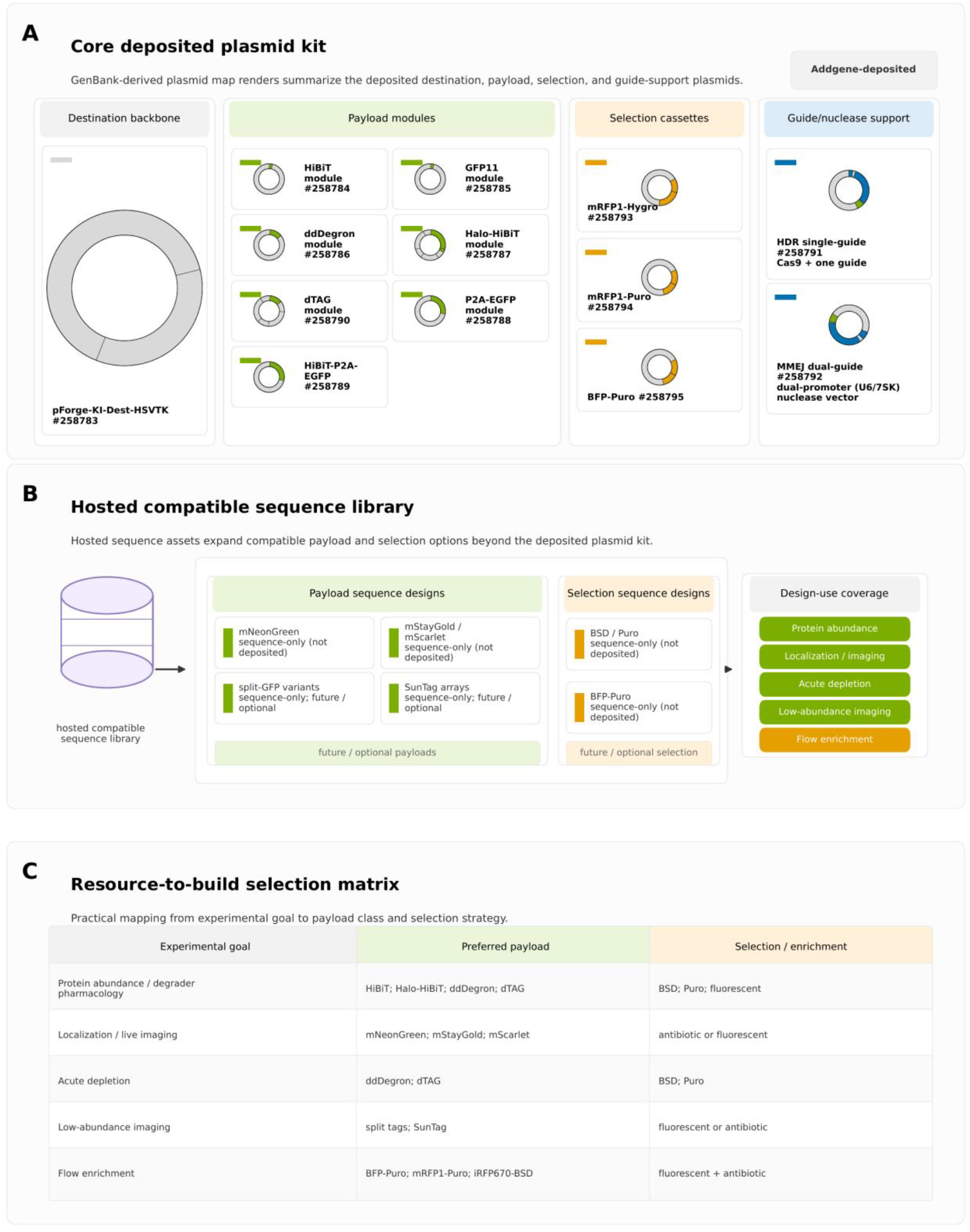
Modular reagent collection and design-use coverage. (A) Core deposited plasmid kit, shown as GenBank-derived plasmid maps: the pForge-KI-Dest-HSVTK destination backbone; reusable payload modules (HiBiT, GFP11, ddDegron, dTAG, Halo-HiBiT, P2A-EGFP, and HiBiT-P2A-EGFP); selectable cassettes (mRFP1-Hygro, mRFP1-Puro, BFP-Puro); and HDR single-guide and MMEJ dual-guide nuclease support. (B) A hosted compatible sequence library extends payload (e.g., mNeonGreen, mStayGold/mScarlet, and repeat-rich split-GFP and SunTag designs that are HDR-only) and selection (BSD/Puro, BFP-Puro) options beyond the deposited kit. (C) Resource-to-build selection matrix mapping experimental goals to preferred payload classes and selection/enrichment strategies.

For large-cargo HDR engineering of the AHR and FOSL1 loci, 5′ and 3′ homology modules were generated and combined with shared fusion, selection, and backbone components, yielding modular donor vectors p1134 (AHR HiBiT-P2A-EGFP) and p1135 (FOSL1 HiBiT-P2A-EGFP). For PITCh/MMEJ-mediated targeting of the IRF1 locus, the entire donor cassette, comprising short microhomology arms, a HiBiT tag, an mNeonGreen reporter, and a loxP-flanked iRFP670-T2A-BSD selection cassette, was synthesized as a single continuous fragment and cloned into an HSV-TK counterselection backbone to generate p1538 (IRF1 MMEJ donor). Locus-specific nuclease guides were adapted to each workflow: single C-terminal guides for AHR and FOSL1 were cloned into a base nuclease vector (p0933), while the IRF1 PITCh pipeline used a dual-guide nuclease vector (p1562) built on the pForge-KI-MMEJ-Cas9-DualGuide architecture to drive both the gene-specific and PITCh-release guides from distinct promoters.

### Assembly of representative HDR and PITCh/MMEJ donors

We assembled endogenous knock-in donor constructs targeting the AHR, FOSL1, and IRF1 loci (Figure 3). Modular HDR donors were generated by combining locus-specific 5′ and 3′ homology arms with the shared HiBiT-P2A-EGFP fusion module, a selectable cassette, and the pForge-KI-Dest-HSVTK acceptor backbone, producing p1134 (AHR), p1135 (FOSL1), and p1130 (IRF1). The representative IRF1 HDR donor p1130 (9,560 bp total) carried 999-bp and 1,010-bp homology arms flanking the HiBiT-P2A-EGFP payload and a BFP-Puro selectable cassette, giving a 5,586-bp functional/integrating segment (Figure 3A). These kilobase-scale arms make single-fragment synthesis of the complete donor impractical, but the multi-part modular framework allowed the locus-specific arms to be customized per locus while reusing identical fusion, selection, and backbone modules.

**Figure 3.**
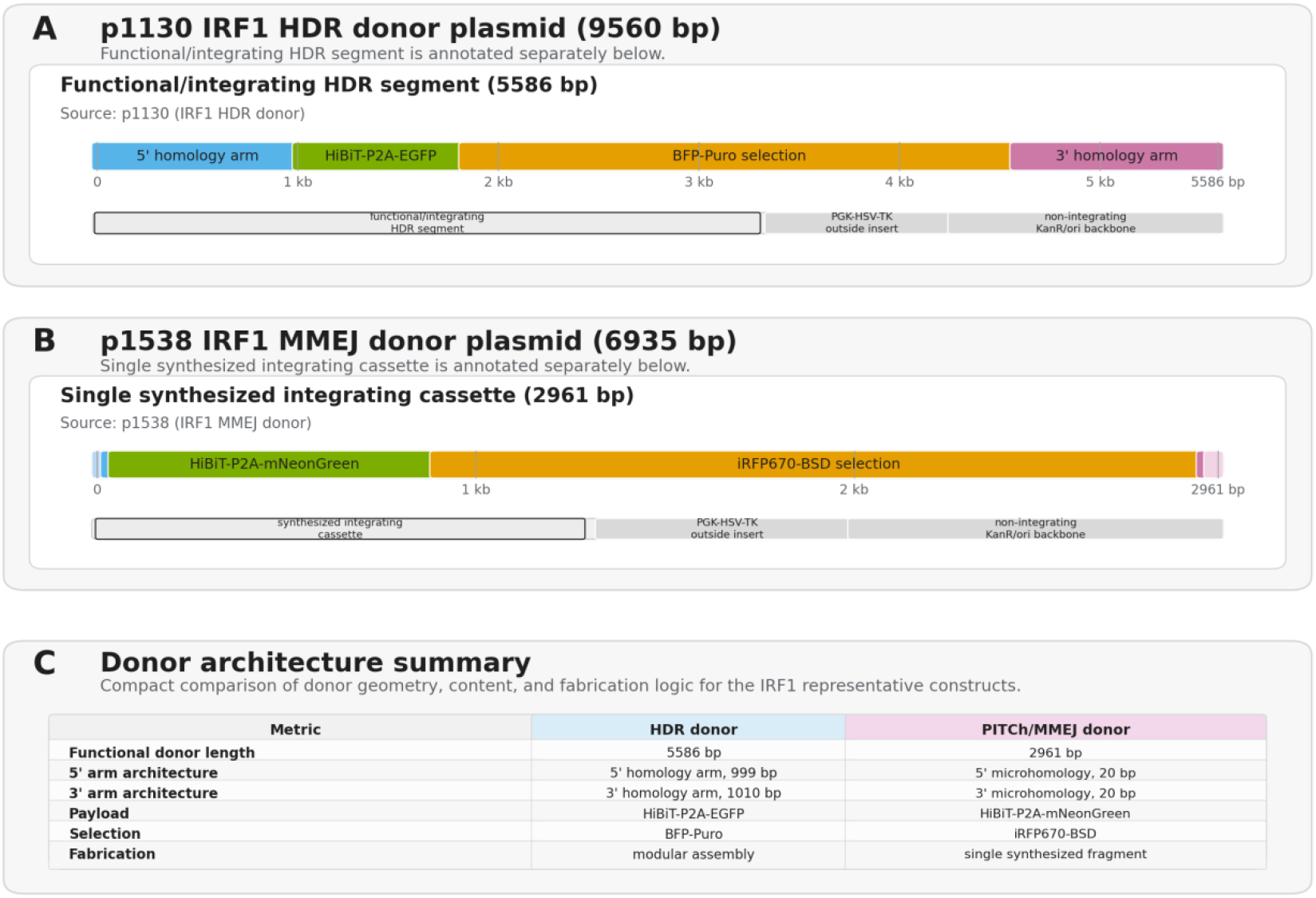
Representative donor architectures, drawn to scale. (A) The IRF1 HDR donor plasmid p1130 (9,560 bp); the 5,586-bp functional/integrating segment comprises a 999-bp 5′ homology arm, the HiBiT-P2A-EGFP payload, a BFP-Puro selection cassette, and a 1,010-bp 3′ homology arm, with PGK-HSV-TK counterselection and a KanR/ori backbone lying outside the integrating segment. (B) The IRF1 PITCh/MMEJ donor plasmid p1538 (6,935 bp); a single synthesized 2,961-bp integrating cassette carries 20-bp microhomology arms, a HiBiT-P2A-mNeonGreen payload, and an iRFP670-T2A-BSD selection cassette, again with PGK-HSV-TK counterselection and a KanR/ori backbone. (C) Quantitative comparison of the two IRF1 donors, highlighting the roughly 50-fold difference in targeting-arm length (approximately 1 kb versus 20 bp) and the modular-assembly vs single-fragment fabrication logic. Feature positions and lengths were parsed from the deposited GenBank files.

For IRF1, single-fragment synthesis additionally yielded the PITCh/MMEJ donor p1538 (6,935 bp total), in which a 2,961-bp integrating cassette containing 20-bp 5′ and 3′ microhomology arms, a HiBiT-P2A-mNeonGreen payload, and a loxP-flanked iRFP670-T2A-BSD selection cassette was synthesized as a single continuous fragment and cloned into a KanR/ori backbone carrying a PGK-driven HSV-TK counterselection marker (Figure 3B,C). To deploy this donor, we engineered p1562 by cloning the IRF1-specific targeting guide into the pForge-KI-MMEJ-Cas9-DualGuide backbone, which expresses the locus guide from a human U6 promoter and the PITCh donor-release guide from a 7SK promoter, a dual-promoter design intended to reduce the repeated-promoter instability associated with some dual-guide lentiviral vectors^18^.

These experiments were not designed as a quantitative comparison of HDR and PITCh/MMEJ editing efficiency. The two workflows differ in cell-line context, payload architecture, selection marker, and validation stage, and we therefore interpret Figure 3 as an operational and architectural comparison of donor fabrication logic rather than a pathway-efficiency benchmark.

### Generation and molecular validation of HDR knock-in reporter pools

To generate endogenous knock-in reporter pools, NCI-H1373 cells were co-transfected with a gene-specific C-terminal CRISPR nuclease (cloned into the p0933 backbone) and the corresponding modular HDR donor using BioT, targeting AHR, FOSL1, and IRF1 for HiBiT-P2A-EGFP insertion (Figure 4A). Three days after transfection, cells underwent puromycin selection followed by recovery, and reporter populations were resolved by flow cytometry on the basis of cassette (BFP) and reporter (GFP) fluorescence; two populations were collected for each target, a BFP-positive population and a BFP/GFP double-positive population. In the representative IRF1 sort, the BFP+/GFP+ fraction was enriched from 0.00% to 11.78% (Figure 4A).

**Figure 4.**
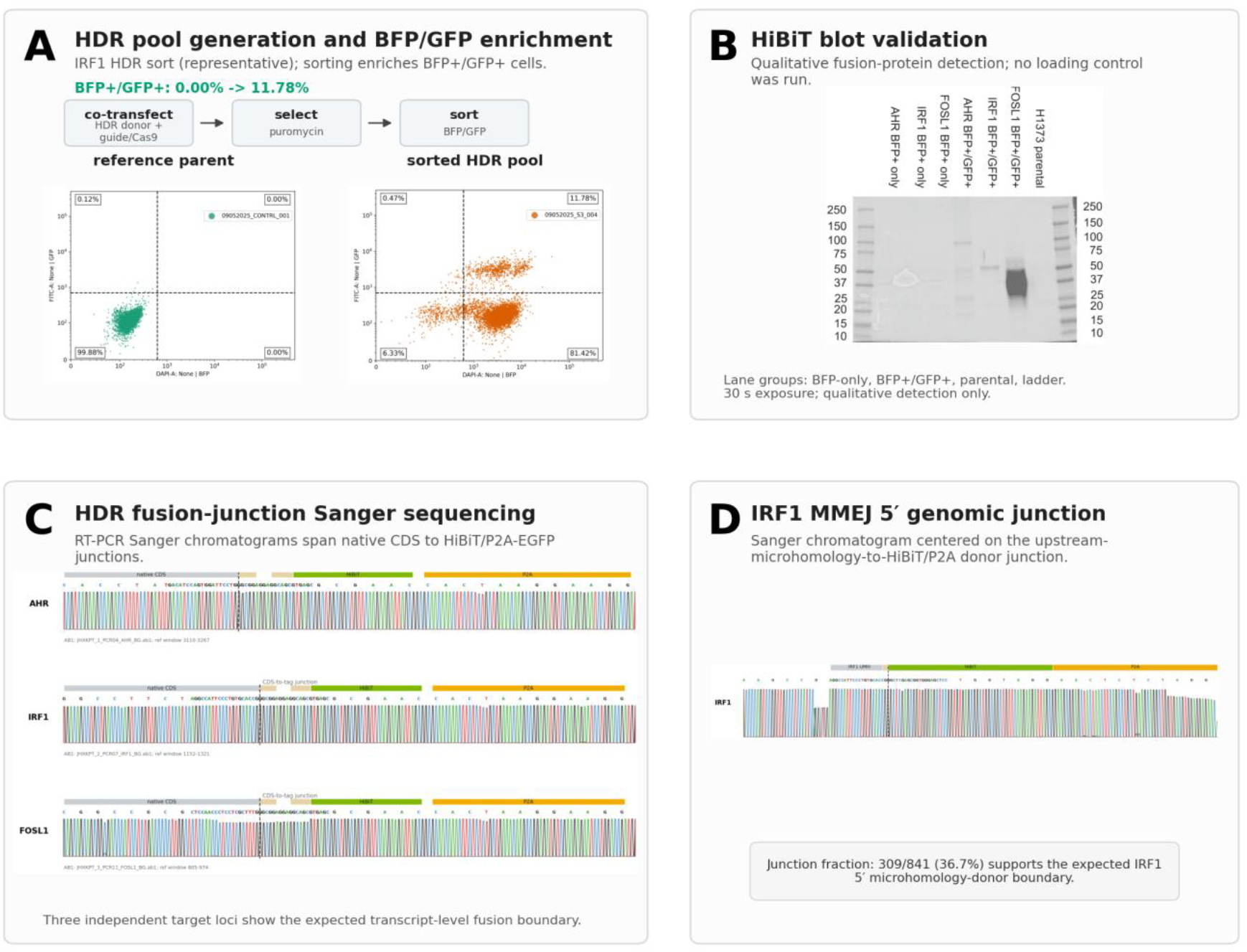
Generation and molecular validation of knock-in reporter pools. (A) HDR pool generation workflow (co-transfection, puromycin selection, and BFP/GFP sorting) and a representative IRF1 sort enriching the BFP+/GFP+ fraction from 0.00% to 11.78%. (B) Nano-Glo HiBiT blot showing fusion-protein signal in the BFP+/GFP+ pools, but not the BFP+-only pools or parental cells, across AHR, IRF1, and FOSL1 (qualitative detection; no loading control; 30-s exposure). (C) RT-PCR Nanopore chromatograms spanning the native CDS–to– HiBiT/P2A-EGFP fusion junction for all three loci. (D) IRF1 PITCh/MMEJ genomic junction: amplicon sequencing recovered the expected 5′ microhomology–donor junction in 309 of 841 reads (36.7%).

For all three loci, the BFP+/GFP+ double-positive population, but not the BFP+/GFP– population, contained detectable HiBiT-tagged protein by Nano-Glo HiBiT blot (qualitative detection; no loading control was run; Figure 4B), indicating that cassette acquisition alone (BFP) was insufficient to mark productive knock-in and that GFP positivity identified cells expressing the endogenous fusion allele. Correct integration was further confirmed at the transcript level: RT-PCR across the fusion junction followed by Nanopore sequencing showed in-frame fusion of the HiBiT-P2A-EGFP cassette at each locus (Figure 4C), with a single silent substitution detected at the FOSL1 junction.

### An inducible, selection-marker-free IRF1 reporter line (K025)

Because IRF1 is an interferon-responsive transcription factor^19^, we asked whether the IRF1 knock-in reporter was cytokine-inducible. Treatment with IFNγ (20 ng/ml, 24 h) increased the BFP+/GFP+ fraction from 47.36% to 73.08% and decreased the BFP+/GFP– fraction from 41.09% to 17.40%, consistent with induction of the endogenous IRF1 reporter (Figure 5A). A weak-basal, inducible subpopulation was isolated by sequential sorting: the high-GFP population showed little further induction, whereas the weak-GFP population was strongly induced by IFNγ plus TNFα, identifying it as the inducible reporter population (Figure 5B).

**Figure 5.**
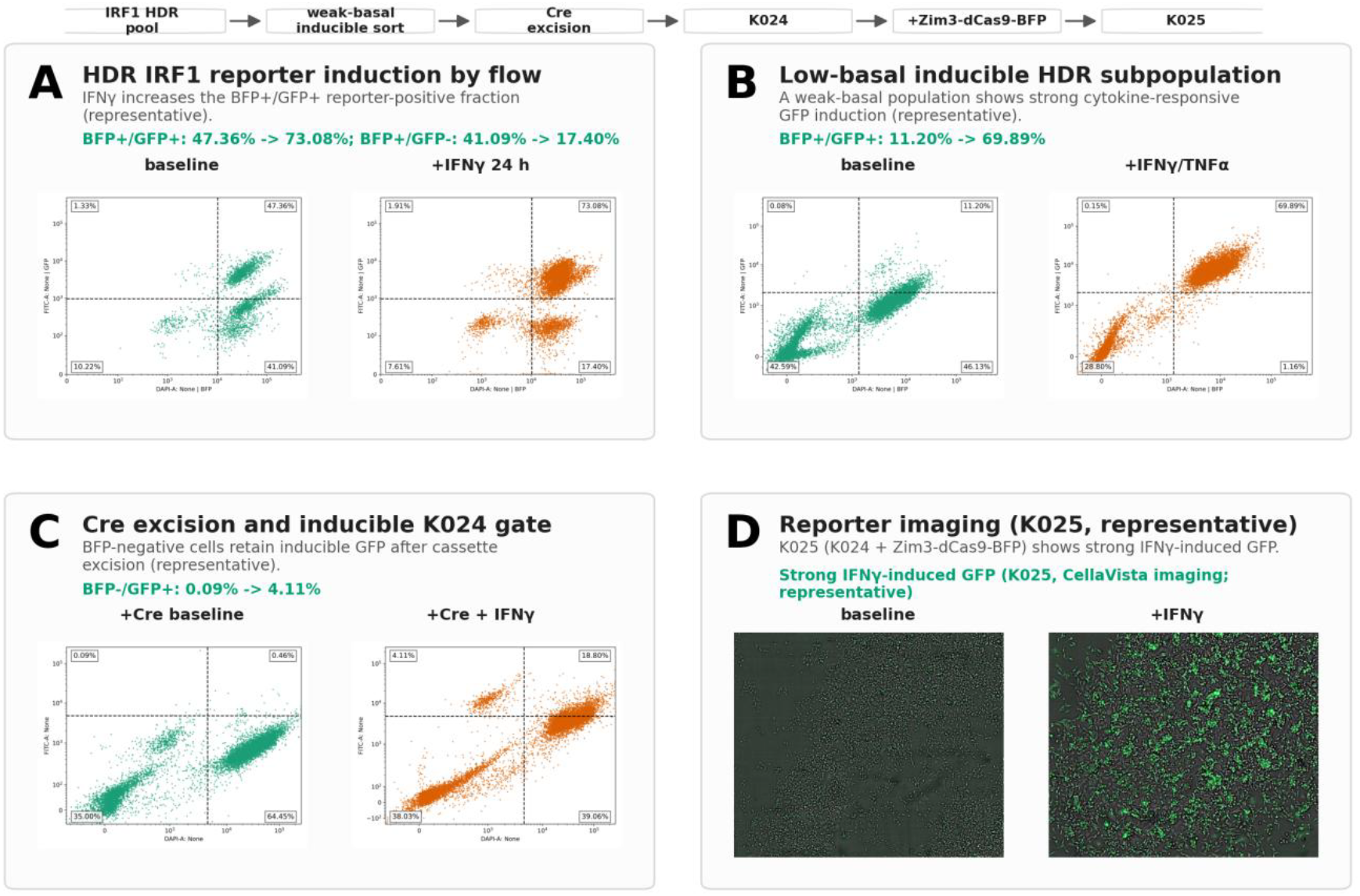
Functional, inducible endogenous IRF1 reporters. (A) IFNγ (20 ng/ml, 24 h) induction of the IRF1 HDR reporter, increasing the BFP+/GFP+ fraction from 47.36% to 73.08% and decreasing the BFP+/GFP– fraction from 41.09% to 17.40%. (B) Sequential sorting isolates a weak-basal subpopulation that is strongly induced by IFNγ plus TNFα, with the BFP+/GFP+ fraction rising from 11.20% to 69.89% (representative dot plots). (C) Cre-mediated removal of the loxP-flanked BFP/puromycin cassette and one-step selection of BFP-negative, inducible cells (K024), with the IFNγ-inducible BFP–/GFP+ fraction rising from 0.09% to 4.11%. (D) K025 (K024 + Zim3-dCas9-BFP) IFNγ induction by CellaVista imaging.

To generate a selection-marker-free reporter line, the loxP-flanked BFP/puromycin cassette was excised by transient Cre, and BFP-negative, IFNγ-inducible cells (4.11%, Figure 5C right panel) were selected in a single step, yielding the inducible reporter line K024. Introduction of Zim3-dCas9-BFP^20^ into K024 produced K025, a Zim3-dCas9–expressing IRF1 reporter line that showed strong IFNγ-induced GFP by CellaVista imaging (Figure 5D), establishing a functional, selection-marker-free, inducible endogenous reporter suitable for downstream CRISPRi perturbation studies.

### Endogenous tagging by single-fragment PITCh/MMEJ

To test the single-fragment PITCh/MMEJ pipeline, NCI-H23 Zim3-dCas9 cells were co-transfected with the synthesized MMEJ donor p1538 (IRF1 HiBiT-P2A-mNeonGreen; loxP-flanked iRFP670-T2A-BSD selection; PGK-HSV-TK counterselection) and the all-in-one dual-guide nuclease p1562. After blasticidin selection, iRFP670+/mNeonGreen+ cells were sorted (K030); because basal mNeonGreen was low, stimulation with IFNγ plus TNFα followed by fluorescence sorting enriched an inducible population, K030i. K030i returned to a low-fluorescence basal state and was re-inducible by IFNγ, demonstrating reversible reporter behavior rather than constitutive expression.

Genomic junction PCR followed by amplicon sequencing confirmed correct integration in a substantial fraction of the pool: of 841 sequenced reads spanning the knock-in junction, 309 (36.7%) carried the expected junction (Figure 4D), consistent with partial enrichment of a polyclonal population^21^. For the HDR pools, productive fusion expression was supported by reporter sorting, HiBiT protein detection, and RT-PCR/Nanopore confirmation of the expressed fusion junction; for the PITCh/MMEJ workflow, pooled genomic amplicon sequencing confirmed enrichment of the expected IRF1 5′ microhomology–donor junction, and cytokine-responsive fluorescence supported functional reporter behavior in the enriched population. These data show that the modular donor architecture supports endogenous tagging by both HDR and PITCh/MMEJ, while clonal allele-level validation remains in progress. Single-cell cloning of K030i to recover monoclonal, junction-confirmed lines is ongoing.

### forgeKI: an automated framework for HDR and PITCh/MMEJ knock-in design

We implemented forgeKI, an R package that automates C-terminal reporter knock-in design across both HDR and PITCh/MMEJ repair pathways (Figure 6). From a target gene symbol and a chosen reporter configuration, forgeKI resolves the locus, transcript, CDS, and terminal stop codon on hg38; enumerates and ranks SpCas9 guides around the C-terminal insertion boundary by PAM, cut geometry, and off-target risk; flags loci that violate standard C-terminal tagging assumptions (for example, selenoproteins, organellar loci, or C-terminal processing and localization motifs); extracts the targeting or microhomology arms with the native stop excluded; proposes Type IIS domestication and Cas9 re-cleavage–blocking edits; validates the resulting virtual edited allele; and constructs route-appropriate donor designs: five-module Golden Gate donors for HDR and single-fragment PITCh donors for MMEJ. It exports synthesis-ready module sequences, guide oligos, annotated GenBank maps, a vendor order sheet, a ranked cell-line shortlist, and a per-design report with ranked guide and donor recommendations. By separating gene-specific design from reusable functional modules, forgeKI mirrors the organization of the physical cloning system. Core donor and guide design are fully reproducible from public genome annotations; the optional cell-line context overlay is provided as a restricted research-use layer and is not required for donor construction.

**Figure 6.**
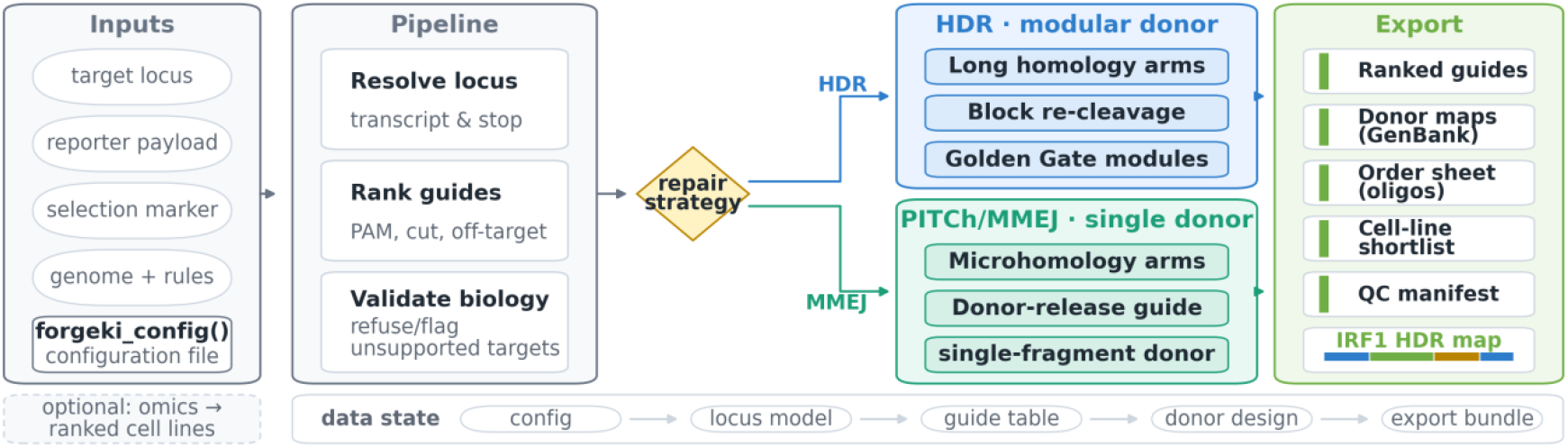
The forgeKI automated design framework. From a single configuration, forgeKI resolves the locus, ranks guides, and validates target biology, then branches by repair strategy to build a complete HDR (modular five-part Golden Gate) or PITCh/MMEJ (single-fragment) donor, and exports a reproducible package: ranked guides, annotated GenBank donor maps, an order sheet, a cell-line shortlist, and a QC manifest. Code and archived releases are listed in Data and Code Availability.

### Summary of evidence and validation status

We use the following terms throughout. Construct validation refers to sequence-confirmed donor plasmids; pooled knock-in validation refers to enriched reporter-positive populations with fusion-junction or protein evidence; functional reporter validation refers to cytokine-responsive reporter behavior; and clonal allele validation refers to monoclonal lines with confirmed 5′ and 3′ junctions and allele structure. The present work reports the first three categories; clonal allele validation is ongoing for selected PITCh/MMEJ-derived lines. Table 1 summarizes the resources, evidence, and current validation status for each construct and pathway.

**Table 1.** Evidence and validation summary across constructs and pathways.

| Construct / resource | Pathway & cell line | Payload | Evidence shown | Status | Limitations / peer-review expansion |
| --- | --- | --- | --- | --- | --- |
| p1134 (+ p0933 nuclease) | HDR · NCI-H1373 | HiBiT-P2A-EGFP | Donor assembled; BFP/GFP-sorted pool; HiBiT blot signal in BFP+/GFP+ pool; RT-PCR/Nanopore fusion transcript | Enriched pool | No genomic/allelic structure (Southern/long-read); not clonal |
| p1135 (+ p0933) | HDR · NCI-H1373 | HiBiT-P2A-EGFP | As AHR; single silent substitution noted at the junction | Enriched pool | No allelic structure; not clonal |
| p1130 (+ p0933); Cre #123133; Zim3-dCas9 #188777 | HDR · NCI-H1373 | HiBiT-P2A-EGFP | Sorted pool; HiBiT; RT-PCR/Nanopore; IFNγ-inducible; K024 after Cre excision; K025 after Zim3-dCas9; IFNγ imaging | Functional reporter line (selection-marker-free) | Not monoclonal; no allele structure; CRISPRi not functionally assayed in K025 |
| p1538 (+ p1562 dual-guide) | PITCH/MMEJ · NCI-H23 Zim3-dCas9 | HiBiT-P2A-mNeonGreen; iRFP670-T2A-BSD | Single-fragment donor; blasticidin selection; iRFP670/mNeonGreen sort (K030); inducible/reversible (K030i); 309/841 reads (36.7%) with expected 5' junction | Inducible enriched pool (proof of principle) | Polyclonal; 5' junction only (no 3'/protein); single-cell cloning in progress |
| forgeKI R package (v0.1.1) | Both pathways (design) | Not applicable | Resolves locus/transcript/CDS/stop; ranks guides; validates biology; builds HDR and PITCH donors; exports guides, GenBank maps, order sheet, cell-line shortlist, QC manifest; unit-tested | Software resource | Off-target assessment required for production guides; per-locus output table not included in this version; cell-line overlay optional/restricted |

## Discussion

We developed a modular framework for endogenous knock-in engineering consisting of reusable fusion modules, selectable cassettes, acceptor backbones, gene-specific targeting elements, and an automated design package. Using this architecture, we generated donor constructs for multiple target genes and implemented both HDR- and PITCh/MMEJ-based integration strategies within a common organizational framework, and we generated a functional, cytokine-inducible IRF1 HDR reporter line together with an inducible IRF1 PITCh/MMEJ reporter pool. Together, these components establish a platform for constructing and deploying endogenous reporter alleles while minimizing redesign of shared donor elements.

We did not aim to define a single optimal knock-in architecture, but to accommodate the diversity of experimental requirements across the field. Quantitative protein measurements may benefit from sensitive tags such as HiBiT, whereas imaging and localization studies often require fluorescent reporters; chemical biology applications may leverage HaloTag-based approaches, and targeted-degradation studies may require degron payloads. Similarly, selection strategies effective for highly expressed genes may be less effective for low-expression genes, motivating independently driven, removable selection cassettes. We therefore sought a framework capable of incorporating multiple validated design strategies rather than prescribing a single implementation.

The execution of both HDR and PITCh/MMEJ targeting illustrates the practical benefit of matching the cloning methodology to the physical scale of the repair mechanism. For microhomology-mediated pathways, minimizing the targeting arms to short approximately 20-bp sequences permits the entire donor payload to be synthesized as a single continuous block, compressing experimental cycle times. For large-cargo conventional HDR insertions, the multi-part modular framework remains the ideal choice to bypass the size limitations of single-piece synthesis. The dual-promoter pForge-KI-MMEJ-Cas9-DualGuide architecture was designed to reduce the repeated-promoter instability associated with some dual-guide vectors, providing a convenient single-vector platform for PITCh applications.

The present study reports enriched reporter pools and a selection-marker-free HDR reporter line rather than monoclonal, allele-validated lines; single-cell cloning of K030i to recover monoclonal, junction-confirmed lines is in progress. Notably, reporter induction was not recovered at every locus attempted (TIPARP, CXCL9, and CXCL10). These failures identify an important design constraint: endogenous reporter engineering is constrained by locus biology and expression state, which is precisely why a pathway-aware and cell-context-aware design framework is needed. Predicting when endogenous expression, cytokine responsiveness, selection architecture, and reporter brightness are jointly sufficient to recover functional knock-in populations is a priority for future development. The optional cell-line context overlay in forgeKI draws on a fixed reference bundle and a restricted-use ranking model and is not required for donor or guide design. Finally, although forgeKI now supports both HDR and PITCh/MMEJ design, broader repair-pathway coverage (for example, HMEJ and HITI) remains an avenue for future development.

In this preprint, we report the reagent collection, the deposited plasmid kit, the software design, and representative biological feasibility. The next phase will convert enriched populations into clonal, allele-validated lines, extend validation to additional loci and cell states, benchmark pathway choice under matched conditions, and expand forgeKI to additional repair architectures.

## Materials and Methods

### Cell lines and culture

HDR-mediated knock-in experiments were performed in NCI-H1373 cells, and MMEJ-mediated knock-in experiments were performed in an NCI-H23-derived cell line stably expressing Zim3-dCas9. NCI-H1373 (ATCC CRL-5866) and NCI-H23 (ATCC CRL-5800) cells were obtained from the American Type Culture Collection. Cells were maintained in RPMI-1640 medium supplemented with 10% fetal bovine serum and penicillin–streptomycin, and were cultured at 37°C in a humidified incubator with 5% CO2. All cell lines tested negative for mycoplasma contamination.

### Stable cell line generation

An NCI-H23 Zim3-dCas9 stable cell line was generated by transducing NCI-H23 cells with concentrated lentivirus encoding Zim3-dCas9-P2A-BFP (pHR-UCOE-EF1a-Zim3-dCas9-P2A-BFP; Addgene #188777), produced by VectorBuilder. Cells were transduced by spinoculation (MOI = 5; 850 × g, 90 min, 32°C), cultured overnight, and transferred to fresh complete medium. Transduced cells were expanded and enriched by two rounds of sorting for BFP-positive cells on a FACSAria III cell sorter (BD Biosciences) at the UCLA JCCC Flow Cytometry Core Facility. Zim3-dCas9 repressor activity was confirmed with the CRISPRiTest Functional dCas9-Repressor Assay Kit (Cellecta) according to the manufacturer’s instructions.

### Modular donor architecture and construction

Knock-in donor constructs were assembled using a five-module architecture consisting of (i) a 5′ targeting arm, (ii) a fusion module, (iii) a selectable cassette, (iv) a 3′ targeting arm, and (v) an acceptor backbone, with gene-specific information localized to the targeting-arm modules. Gene-specific targeting modules were designed as synthetic DNA fragments (Twist Bioscience) carrying standardized Type IIS assembly overhangs compatible with the Mobius Universal Assembly Vector (mUAV) system; overhangs were selected using optimized assembly junctions^22^. Fusion, selectable-cassette, and gene-specific modules were cloned into mUAV entry vectors using AarI- or PaqCI-mediated Golden Gate assembly^23^.

Complete HDR donor constructs were assembled by multi-part BsaI Golden Gate cloning, combining individual mUAV entry modules^24^ with the pForge-KI-Dest-HSVTK destination vector. The PITCh/MMEJ donor p1538 was fabricated in a single step by synthesizing the entire cassette, comprising microhomology targeting sequences, HiBiT tag, mNeonGreen reporter, loxP-flanked iRFP670-T2A-BSD selection cassette, and PGK-driven HSV-TK counterselection marker, as a single continuous fragment (Twist Bioscience), followed by direct cloning into a stable KanR/ori bacterial backbone. All assembly products were transformed into E. coli, screened by restriction digest and sequencing, and propagated for downstream use.

### HDR and PITCh/MMEJ donor and guide design

HDR donors carried locus-specific 5′ and 3′ homology arms on the kilobase scale (999 bp and 1,010 bp in the representative IRF1 donor p1130) flanking the fusion and selectable-cassette modules. PITCh/MMEJ donors used short locus-specific approximately 20-bp microhomology arms flanking the payload, together with flanking PITCh donor-release guide sequences. Single guides for HDR were cloned into nuclease vectors derived from p0933 (pForge-KI-HDR-Cas9-SingleGuide). For PITCh/MMEJ, the all-in-one nuclease vector p1562 was generated by cloning the IRF1-locus targeting guide into the pForge-KI-MMEJ-Cas9-DualGuide backbone, which drives the gene-specific guide from a human U6 promoter and the PITCh donor-release guide from a 7SK promoter.

### Generation of knock-in cell pools

Knock-in cell pools were generated by either homology-directed repair (HDR) or microhomology-mediated end joining (MMEJ). For both methods, cells were seeded at 300,000 cells per well in 6-well plates and co-transfected with 2 μg total donor and nuclease plasmid DNA using 3 μl BioT reagent per well according to the manufacturer’s instructions, with six wells transfected per experiment. For HDR-mediated knock-in, NCI-H1373 cells were selected with puromycin (1.5 μg/ml) beginning 3 days after transfection for 4 days, then expanded and enriched by flow sorting for BFP and GFP expression. Where indicated, the loxP-flanked puromycin–BFP cassette was excised by transient transfection of pCMV-Cre (Addgene #123133), followed by sorting for BFP-negative cells to generate K024. K024 cells were then transduced with high-titer lentiviral Zim3-dCas9-P2A-BFP (Addgene #188777; VectorBuilder) and enriched by two rounds of BFP-positive sorting to generate K025. For MMEJ-mediated knock-in, NCI-H23 Zim3-dCas9 cells were selected with blasticidin (6 μg/ml) beginning 3 days after transfection for 7 days, then expanded and enriched by flow sorting for iRFP670 (APC) and mNeonGreen (FITC) expression. All sorting was performed on a FACSAria III (BD Biosciences) at the UCLA JCCC Flow Cytometry Core Facility. Negative selection against random integration via the HSV-TK marker was not applied in the present workflow; enrichment relied on antibiotic selection and fluorescence sorting.

### Cytokine induction

For cytokine induction, cells were treated with recombinant human IFNγ or a combination of IFNγ and TNFα as indicated. For IFNγ-only stimulation, cells were treated with IFNγ at 20 ng/ml for 24 h (R&D Systems, 285-IF-100). For combined stimulation, cells were treated with IFNγ at 10 ng/ml and TNFα at 10 ng/ml for 24 h (R&D Systems, 210-TA-020). Following treatment, cells were analyzed by flow cytometry as described above or by CellaVista imaging (SynenTec).

### Junction PCR and amplicon sequencing

To detect the upstream fusion junctions in AHR, IRF1, and FOSL1 mRNAs from NCI-H1373 knock-in pools, total RNA was isolated using the RNeasy Mini Kit (Qiagen) and cDNA was synthesized by reverse transcription with the MultiScribe Reverse Transcriptase Kit (Thermo Fisher Scientific). To detect the upstream junction in IRF1 genomic DNA from NCI-H23 Zim3-dCas9 knock-in pools, genomic DNA was isolated using the PureLink Genomic DNA Mini Kit (Invitrogen). For PCR amplification, 25 ng of cDNA or genomic DNA was used as template in a 25 μl reaction containing 1× Phusion HF buffer, 200 μM dNTPs, 500 nM each forward and reverse primer, and 0.5 U Phusion High-Fidelity DNA Polymerase (Thermo Scientific, F530). Cycling conditions were 98°C for 30 s; 30 cycles of 98°C for 10 s, 55°C for 30 s, and 72°C for 30 s; and a final extension at 72°C for 10 min. Products were resolved on a 2% agarose gel, and bands of the expected size were excised and purified with the NucleoSpin Gel and PCR Clean-Up Kit (Takara Bio). HDR fusion-junction amplicons (AHR, IRF1, and FOSL1 cDNA) were Nanopore-sequenced (Figure 4C), whereas the IRF1 MMEJ genomic-junction amplicon was sequenced on the Oxford Nanopore platform (Plasmidsaurus), with 309 of 841 junction-spanning reads (36.7%) carrying the expected sequence (Figure 4D). Primer sequences are listed in Table 2 (HDR) and Table 3 (MMEJ).

**Table 2.**
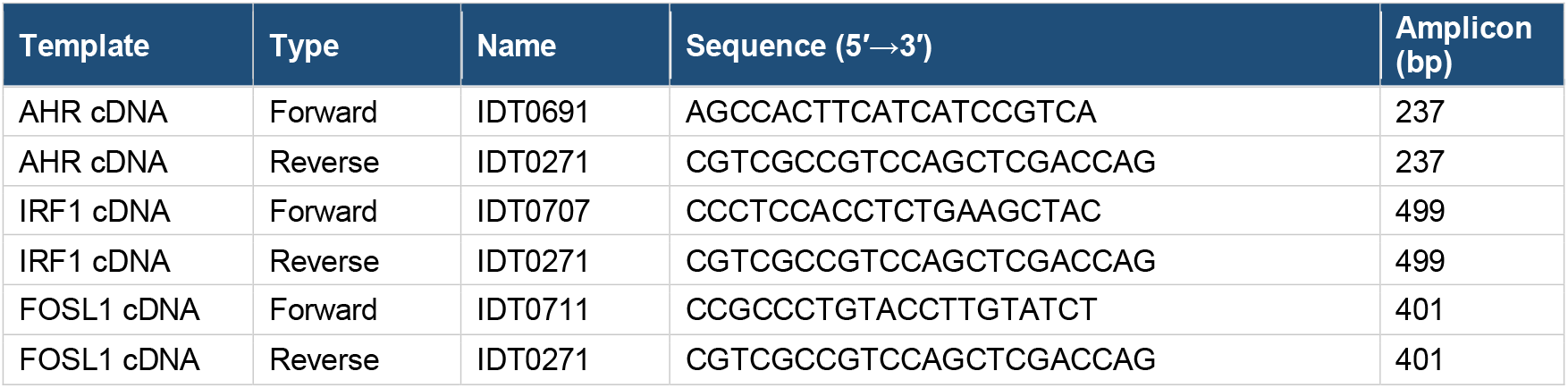
Primers for HDR upstream-junction amplification.

**Table 3.**
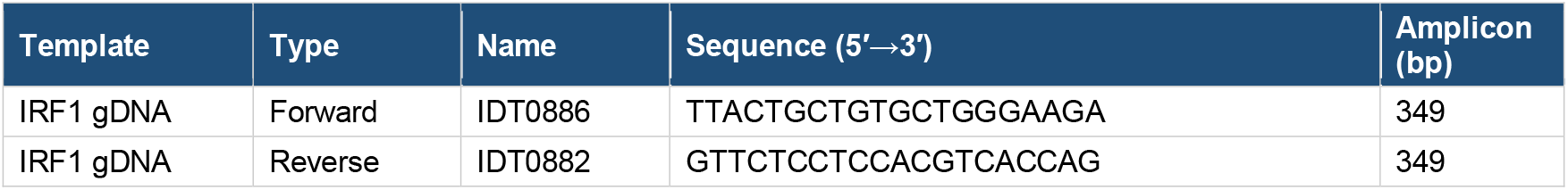
Primers for MMEJ upstream-junction amplification.

### Nano-Glo HiBiT protein detection

For HiBiT protein detection, cells were lysed in protein lysis buffer supplemented with protease and phosphatase inhibitors, and protein concentration was determined by BCA assay. Equal amounts of protein lysate (typically 20 μg per sample) were resolved by SDS-PAGE and transferred to PVDF membranes. HiBiT-tagged proteins were detected using the Nano-Glo HiBiT Blotting System (Promega) according to the manufacturer’s protocol, and luminescent signal was captured on a ChemiDoc imaging system (Bio-Rad). Detection was qualitative, and no loading control was run.

### Computational design framework (forgeKI)

Donor design was automated with forgeKI, an R package implementing a staged, repair-pathway-aware knock-in design pipeline for both HDR and PITCh/MMEJ. Configuration is specified through explicit, serializable option objects (gene, transcript override, guide search radius, homology-arm lengths, repair strategy, Type IIS/domestication policy, Golden Gate overhangs, off-target mode, and optional cell-line reference), and each run executes in an isolated job directory with structured logging and per-stage outputs. The pipeline resolves the locus, transcript, CDS, and terminal stop codon and computes the C-terminal insertion coordinate immediately upstream of the native stop (hg38 via Bioconductor BSgenome/TxDb resources, MANE-preferred transcript selection); enumerates and annotates SpCas9 guides around the insertion boundary (PAM, cut geometry, GC, homopolymer, and U6/polyT); assesses off-target risk (modes none, auto, exact_genome, exact_hg38); and validates target biology, refusing or flagging loci that violate standard C-terminal tagging assumptions. It then branches by repair strategy: for HDR, transcript-oriented homology-arm extraction with native-stop exclusion and salvage-length tiers, biology-aware Type IIS domestication, recursive Cas9 re-cleavage blocking, virtual edited-allele validation, and five-module Golden Gate donor construction via a module registry; for PITCh/MMEJ, microhomology-arm design, donor-release-guide handling, virtual-junction validation, and single-fragment donor construction. Design scoring and primary-recommendation logic, and an optional cell-line context overlay drawn from a fixed reference bundle, complete the run. Off-target assessment is required for a production recommendation; runs executed with off-target mode “none” are used for donor-construction validation only and are not promoted as production-ready guide selections. forgeKI depends on standard Bioconductor genome/annotation packages and is unit-tested across stages.

## Data and Code Availability

### Code availability

The forgeKI knock-in design package is available at https://github.com/dylanconklin-cpu/forgeKI and archived at Zenodo (v0.1.1; https://doi.org/10.5281/zenodo.20683177), released under the MIT license. forgeKI automates HDR and PITCh/MMEJ donor design as described; the cell-line context layer operates as a read-only consumer of a fixed reference bundle and does not include or redistribute the underlying DepMap/CCLE/RRBS-derived feature data or the optional, restricted-use cell-line ranking model, which is not required for donor or guide design. The archived forgeKI release (v0.1.1) includes the PITCh/MMEJ design path and the target-biology validation layer. The cell-line context reference bundle is archived separately at Zenodo (v0.1.0; https://doi.org/10.5281/zenodo.20680156) under restricted research-use terms consistent with DepMap/Broad/CCLE licensing.

### Data availability

Plasmids are deposited at Addgene (deposit 87812); internal construct identifiers, deposited plasmid names, and Addgene catalog numbers are listed in Table 4. Annotated plasmid sequences are provided as Supplementary GenBank files.

**Table 4.**
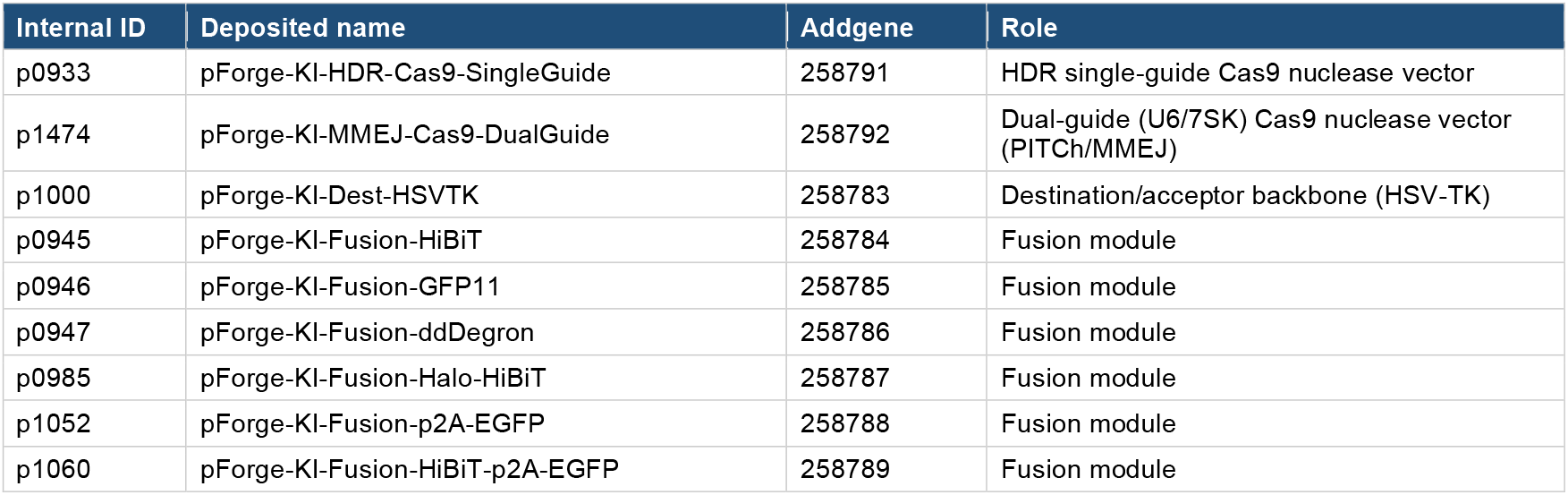
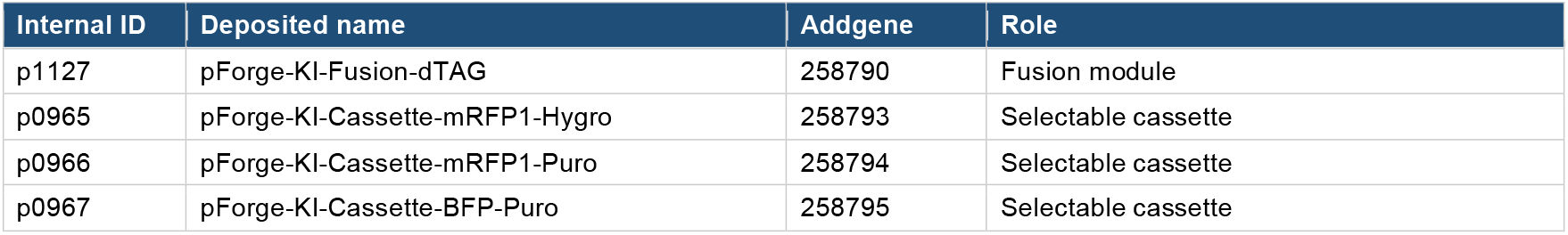
Deposited plasmid kit (Addgene deposit 87812). Internal construct identifiers map to deposited plasmid names and Addgene catalog numbers.

## Author Contributions

D.C., J.-A.L., and M.P. contributed equally to this work. Conceptualization, D.C., J.-A.L., and M.P.; methodology, D.C., J.-A.L., and M.P.; software, D.C.; investigation, D.C., J.-A.L., and M.P.; formal analysis, D.C., J.-A.L., and M.P.; original-draft preparation, D.C., J.-A.L., and M.P.; manuscript review and editing, all authors; supervision, S.M.D. and J.M.L.; resources, S.M.D. and J.M.L.; funding acquisition, S.M.D. and J.M.L. All authors read and approved the final manuscript.

## Acknowledgments

Flow cytometry was performed in the UCLA Jonsson Comprehensive Cancer Center (JCCC) and Center for AIDS Research Flow Cytometry Core Facility, supported by National Institutes of Health awards P30 CA016042 and 5P30 AI028697, and by the JCCC, the UCLA AIDS Institute, the David Geffen School of Medicine at UCLA, the UCLA Chancellor’s Office, and the UCLA Vice Chancellor’s Office of Research.

